# Lentiviral-mediated gene complementation rescues pathogenic *ABCA3* variants

**DOI:** 10.1101/2025.08.29.673150

**Authors:** Ashley L. Cooney, Shakayla Lamer, Ping Yang, Daniel J. Wegner, Frances V. White, F. Sessions Cole, Chris Wohlford-Lenane, Erin Hennessey, Pushpinder Bawa, Darrell N. Kotton, Patrick L. Sinn, Jennifer A. Wambach, Paul B. McCray

**Affiliations:** University of Iowa, Stead Family Department of Pediatrics, Iowa City, IA, 52245, USA; University of Iowa, Center for Precision Medicine for Cystic Fibrosis, Iowa City, IA, 52245, USA; Pappajohn Biomedical Institute, Iowa City, IA, 52245, USA; Center for Regenerative Medicine of Boston University and Boston Medical Center, Boston, MA 02118, USA; Edward Mallinckrodt Department of Pediatrics, Washington University in St. Louis, St. Louis Children’s Hospital, St. Louis, MO 63110, USA

**Keywords:** ABCA3 deficiency, gene therapy, lentiviral vectors, surfactant dysfunction, lamellar bodies, alveolar epithelial type 2 cells

## Abstract

The ATP-binding cassette subfamily A member 3 (ABCA3) protein on the limiting membrane of lamellar bodies in alveolar type 2 (AT2) cells transports phospholipids required for pulmonary surfactant assembly. ABCA3 deficiency results from biallelic pathogenic variants in *ABCA3* and causes progressive neonatal respiratory failure or childhood interstitial lung disease (chILD). Supportive/compassionate care or lung transplantation are the only current definitive treatments for ABCA3 deficiency and progressive respiratory failure. Complementing dysfunctional ABCA3 by gene addition has therapeutic potential. Previous studies show that repairing or complementing *ABCA3* in induced pluripotent stem cell (iPSC)-derived AT2 cells rescues lamellar body morphology and surfactant phospholipid composition. Pathogenic variants disrupt ABCA3 function through altered protein trafficking (type 1) or by impaired phospholipid transport (type 2) into lamellar bodies. Here we tested *ABCA3* gene complementation using a human pulmonary epithelial cell line (A549) with a genomically silenced *ABCA3* locus (*ABCA3*^KO^). From this line, additional cell lines that stably express individual *ABCA3* variant cDNA constructs from a single genomic locus were tested: L101P (type 1), E292V (type 2), E690K (type 2), or wild-type *ABCA3*. Lentiviral-mediated *ABCA3* delivery to each cell line partially rescued localization to LAMP3+ vesicles, lamellar body-like structure morphology, and cell proliferation. A functional assay measuring NF-κB signaling suggested that *ABCA3* complementation ameliorated aberrant inflammatory signaling in E292V or E690K (type 2) mutant lines, but not in L101P (type 1) or knockout lines. These studies highlight the therapeutic potential of gene addition as well as differences between *ABCA3* pathogenic variants that may influence genetic therapy outcomes.

## INTRODUCTION

ATP-binding cassette subfamily A member 3 (ABCA3) deficiency is the most common genetic cause of surfactant dysfunction. Biallelic frameshift or nonsense *ABCA3* variants lead to neonatal respiratory failure and death within the first year of life without lung transplant^1–4^. The respiratory phenotypes and disease outcomes for individuals with biallelic *ABCA3* missense variants are variable and include neonatal respiratory failure, childhood interstitial lung disease (chILD), or later-onset pulmonary fibrosis^1,2^. There is a pressing need to advance new therapies for infants and children with respiratory failure due to ABCA3 deficiency. Restoring surfactant function with a gene complementation strategy is a variant agnostic approach. However, whether complementing ABCA3 will rescue disease phenotypes in the presence of a missense or a loss-of-function *ABCA3* variant remains unknown. Here, we used a human pulmonary epithelial cell line (A549) model to investigate the feasibility of lentiviral (LV) mediated *ABCA3* gene addition for rescuing several functional endpoints related to ABCA3 deficiency.

Functionally, ABCA3 is a phospholipid transporter located in the outer limiting membrane of lysosomally-derived lamellar bodies in alveolar epithelial type 2 (AT2) cells and is critical for surfactant production and function^5^. The ATP hydrolysis activity of ABCA3 plays an important role in phospholipid transport and lamellar body formation^6–8^. The ∼5.2 kb *ABCA3* cDNA encodes a 1,704 amino acid polypeptide which is post-translationally modified before it trafficks to lamellar bodies. N-terminal glycosylation creates a full-length protein with a molecular weight of ∼190 kDa. Subsequent cleavage by cathepsin-L trims the protein to ∼150 kDa^9,10,11^. Wild-type lamellar bodies are distinct, loosely packed, concentric, multilamellar structures, while lamellar bodies from infants and children with ABCA3 deficiency are smaller and electron dense, amorphous or with tightly packed concentric membranes, with some imparting a “fried egg” appearance^12–15^.

Two mechanistically distinct classes of pathogenic *ABCA3* missense variants have been described^16–18^. Type 1 mutants (e.g., L101P) disrupt intracellular trafficking and processing with retention of the mutant protein in the endoplasmic reticulum (ER). In experimental models using an A549 cell line with endogenous *ABCA3* expression genetically silenced, stable expression of the *ABCA3* L101P (type 1, mistrafficking) variant resulted in ER retention of the mutant protein, impaired protein processing, and smaller lamellar body-like structures, similar to those seen among infants and children with ABCA3 deficiency^17,19^. Type 2 mutants (e.g., E292V, E690K) traffic to lamellar bodies but fail to transport surfactant phospholipids due to impaired ATP hydrolysis activity^20^. Both ABCA3 mutant types result in smaller lamellar bodies and alter surfactant phospholipid composition and function^14,15,20–23^.

Studies in cell models suggest several adverse consequences of *ABCA3* variants beyond altered surfactant. Sun et al. recently generated iPSC-derived AT2 (iAT2) cells from infants homozygous for pathogenic *ABCA3* type 2 variants W308R or E690K^20^. These iAT2 cells demonstrated physiologically relevant features, including reduced lamellar body size and altered composition of surfactant phospholipids. Compared to their syngeneic gene edited, corrected controls, mutant iAT2^W308R^ and iAT2^E690K^ cells also demonstrated diminished progenitor potential, increased NF-κB signaling, and pro-inflammatory cytokine production. These results suggest that epithelial-intrinsic aberrant phenotypes are associated with pathogenic *ABCA3* variants. Following tamoxifen induced gene deletion in *Abca3* conditional knockout mice, AT2 cells with residual *ABCA3* expression expanded, and those mice recovered from respiratory failure. These results suggest that corrected AT2 cells may have a proliferative advantage^24^. Complementing ABCA3 deficiency with gene addition might similarly confer a survival/proliferative advantage to treated AT2 cells.

We hypothesized that gene addition of a full length, wild-type *ABCA3* cDNA to A549 cells that stably express individual *ABCA3* pathogenic variants will rescue cellular phenotypic characteristics associated with ABCA3 deficiency, regardless of the disease causing *ABCA3* variant^20^. To date, the impact of complementing wild-type *ABCA3* across different variants in the presence of mutant ABCA3 protein has not been explored. We investigate the utility of a lentiviral vector that integrates into the host genome for prolonged expression and that could potentially last the lifetime of the patient^25^. We use an HIV-based vector expressing wild-type *ABCA3* (5.2 kb) to transduce A549 cells that each stably express one of three disease-causing *ABCA3* variants: L101P, E292V, or E690K, as well as *ABCA3* knockout cells. We report rescued protein processing, increased diameter of ABCA3+ vesicles, and improved lamellar body-like morphology. Gene addition also rescued NF-κB-mediated pro-inflammatory irregularities in cells with phospholipid transport (type 2) mutants E292V and E690K, but not in cells that stably express the mistrafficking (type 1) L101P mutant. However, in both mutant types, increased cell proliferation in complemented cells suggests that this approach may confer a selective advantage to corrected cells. Results from this study are an important step to advance gene therapy for ABCA3 deficiency.

## RESULTS

### A549 cells that express ABCA3 pathogenic variants

We used “landing pad” technology to generate A549/*ABCA3*^KO^ cells as previously described^17^. Briefly, using genome editing, biallelic frameshift mutations introduced into exon 5 to abrogate endogenous *ABCA3* activity, termed *ABCA3^KO^* (**Fig. 1A**). Next, a LV vector containing a CMV promoter driving a LoxFAS-GFP-LoxP cassette (the “landing pad”) was delivered to *ABCA3^KO^*A549 cells, and cells were sorted for GFP expression. Through dilutional cloning and inverse PCR, individual clones were selected to develop a parental cell line with intergenic insertion of the landing pad on chromosome 12. As indicated in **Fig. 1A** step 5, LoxFAS-ABCA3:mCherry-LoxP cassettes carrying L101P, E292V, E690K, or wild-type *ABCA3* were co-transfected with Cre-recombinase for targeted insertion at the landing pad. Successful targeting results in mCherry^+^/GFP^-^ cells that allow for positive and negative selection. The resultant 5 cell lines include: ABCA3^KO^, ABCA3^L101P^, ABCA3^E292K^, ABCA3^E690K^, and ABCA3^WT^. Except for the ABCA3^KO^ cells, each cell line expresses ABCA3 with a C-terminal mCherry tag to visualize mutant ABCA3 trafficking.

**Figure 1:**
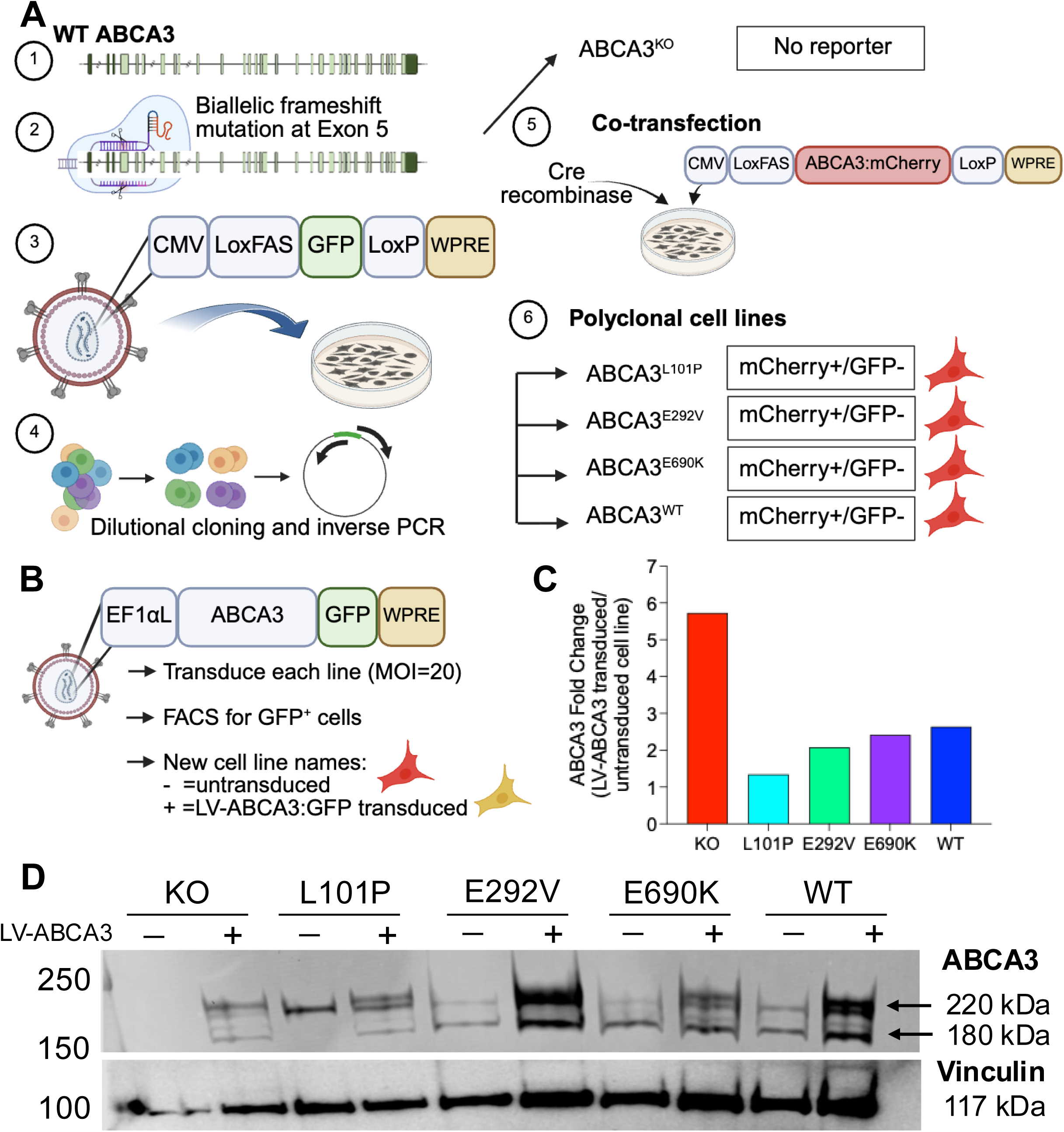
Lentiviral-mediated ABCA3 delivery to A549 cell variants. A) A549/ABCA3 knockout (KO) cells were modified to stably express pathogenic *ABCA3* variants through a series of steps: 1) Exon schematic of wild-type (WT) *ABCA3*. 2) To generate A549/ABCA3^KO^ (knockout) cells, CRISPR/Cas9 was used to introduce a frameshift variant at exon 5. 3) A lentiviral vector carrying CMV-LosFAS-GFP-LoxP was delivered to A549 ABCA3^KO^ cells to create a “landing pad” for stable insertion of *ABCA3* variants at a single genomic locus. 4) GFP positive cells were selected, diluted as single clones, and expanded. 5) Cre-recombinase and LoxFAS-CMV-ABCA3:mCherry-LoxP (and L101P, E292V, E690K variants) were co-transfected into A549 ABCA3^KO^ cells. 6) A total of 5 polyclonal cell lines were generated as outlined. B) A lentiviral vector carrying EF1⍺L-ABCA3:GFP was delivered to all 5 cell lines (MOI=20) and cells were sorted by flow cytometry for GFP positive cells. Throughout the manuscript, a “-” indicates the untransduced, parental cell line and a “+” indicates a ABCA3:GFP^+^ sorted cell line. C) *ABCA3* transcripts were detected by qRT-PCR. Fold change presented as LV-ABCA3:GFP transduced cells/untreated parental lines. D) Immunoblotting results show detection of ABCA3 bands at 220 kDa (uncleaved form) and 180 kDa (post-cleavage form). Vinculin was used as a loading control.

### Lentiviral mediated complementation of ABCA3-deficiency

To determine if *ABCA3* complementation rescues ABCA3 protein expression and function in both mistrafficking (type 1) and phospholipid transport (type 2) mutants, we employed our previously reported VSVG pseudotyped HIV-based LV vector that encodes an ABCA3:GFP cassette driven by a constitutively and ubiquitously active EF1αL promoter (hereafter “LV-ABCA3:GFP”, **Fig. 1B**)^20^. Each of the 5 cell lines was transduced with LV-ABCA3:GFP (MOI=20), resulting in constitutive expression of the ABCA3:GFP fusion protein (**Fig. 1B**). Four days later, GFP^+^ cells were isolated by two rounds of fluorescence activated cell sorting (FACS) (**Supplemental Fig. 1A, B**). In total, 10 cell lines were generated (**Table 1**). Using qRT-PCR, we measured *ABCA3* mRNA levels and observed that the fold change of LV-ABCA3:GFP transduction was similar to wild-type for each cell line that expresses a pathogenic *ABCA3* variant (**Fig. 1C**). As expected, ABCA3^KO^ cells had the largest change in expression following LV correction due to low or undetectable baseline *ABCA3* mRNA levels.

**Table 1:** Reporter genes expressed in each cell line.

|  | Cell line | Reporter genes expressed |
| --- | --- | --- |
| 1 | KO | None |
| 2 | KO + LV-ABCA3:GFP | mCherry<sup>-</sup>/GFP<sup>+</sup> |
| 3 | L101P | mCherry<sup>+</sup>/GFP<sup>-</sup> |
| 4 | L101P + LV-ABCA3:GFP | mCherry<sup>+</sup>/GFP<sup>+</sup> |
| 5 | E292V | mCherry<sup>+</sup>/GFP<sup>-</sup> |
| 6 | E292V + LV-ABCA3:GFP | mCherry<sup>+</sup>/GFP<sup>+</sup> |
| 7 | E690K | mCherry<sup>+</sup>/GFP<sup>-</sup> |
| 8 | E690K + LV-ABCA3:GFP | mCherry<sup>+</sup>/GFP<sup>+</sup> |
| 9 | WT | mCherry<sup>+</sup>/GFP<sup>-</sup> |
| 10 | WT + LV-ABCA3:GFP | mCherry<sup>+</sup>/GFP<sup>+</sup> |

To characterize expression of the ABCA3:GFP fusion proteins from all cell lines (expected sizes: 220 kDa (ABCA3 uncleaved+GFP), 180 kDa (ABCA3 cleaved+GFP)), we performed a western blot using an anti-ABCA3 antibody. We observed increased protein expression in the cells transduced with LV-ABCA3:GFP (**Fig. 1D**). In untransduced cells that express E292V, E690K, or WT ABCA3, we observed both the 220 kDa and 180 kDa bands. This was expected because WT and type 2 variants undergo post-Golgi processing/cleavage^17^. Untransduced ABCA3^KO^ and ABCA3^L101P^ cells lack the 180 kDa band due to retention in the ER and failure to undergo proteolytic cleavage. In summary, LV-mediated gene complementation restored ABCA3 protein expression in each A549 variant cell line, including the fully processed 180 kDa band in ABCA3^KO^ and ABCA3^L101P^ cells, suggesting that complemented ABCA3 produces a mature protein in both type 1 and type 2 ABCA3 mutant backgrounds.

We performed bulk mRNA sequencing to profile the transcriptional landscape in the engineered cell lines (5 cell lines total). Pairwise comparisons of differentially expressed genes (DEGs) between KO and *ABCA3* variant cell lines were performed using gene set enrichment analysis (GSEA) utilizing the MSigDM Hallmark gene sets. We found enrichment for signatures associated with cell proliferation (E2F targets, G2M checkpoint, Myc targets v1 and v2, and mitotic spindle). In addition, reductions in signatures associated with interferon responses relative to ABCA3^KO^ cells were observed for each condition except L101P (**Fig. 2A-D**). Consistent with previous reports in iAT2 cells, loss of functional ABCA3 leads to reduced cell proliferation^20^.

**Figure 2:**
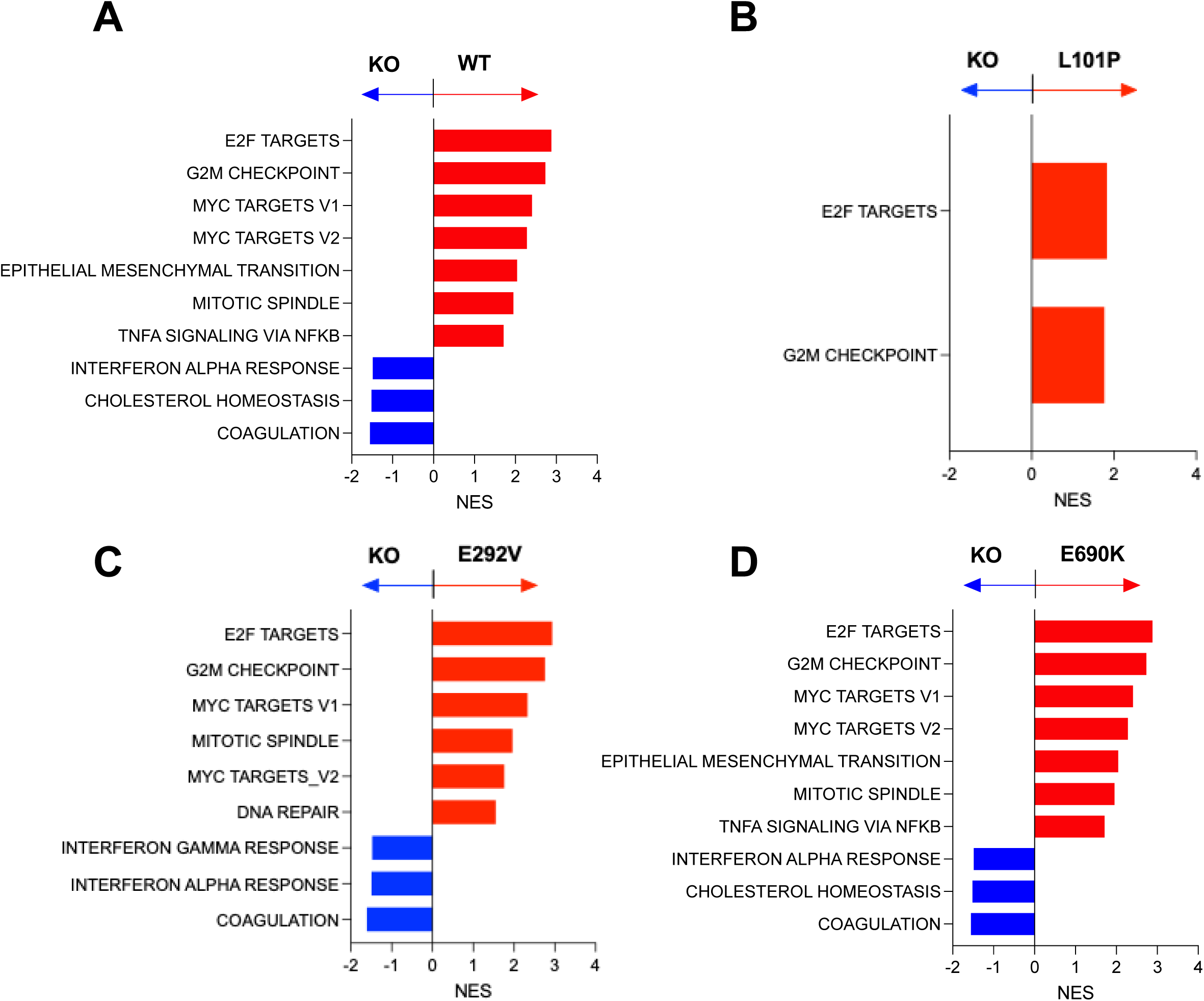
Bulk RNA sequencing of A549 ABCA3 cells. For each cell line carrying wild-type or ABCA3 pathogenic variants, differential expression analysis was compared to ABCA3^KO^ cells. Gene expression was ranked and analyzed by GSEA using the Molecular Signatures Database (MSigDB) Hallmark Gene sets. In each panel, GSEA output represents results for the variant lines vs. knockout. A-D) Bar graphs show normalized enrichment score (NES) for false discover rate (FDR) <0.05. Genes were ranked by log-fold change and tested for enrichment using MSigDB Hallmark gene sets. Pathways increased upon relative to KO are shown in red (positive NES), and those decreased are shown in blue (negative NES). N=4 replicates.

To assess cellular localization of LV-derived ABCA3, we used confocal fluorescence microscopy to localize GFP in the ABCA3:GFP fusion protein or mCherry in the ABCA3:mCherry fusion protein from each cell line. As expected, no GFP or mCherry fluorescence was observed in untransduced ABCA3^KO^ cells (**Fig. 3A**). Since ABCA3^WT^ cells were engineered to include wild-type ABCA3:mCherry at the landing pad site, after transduction of LV-ABCA3:GFP, all GFP tagged transgene expression overlapped with wild-type ABCA3:mCherry. These cells expressed mCherry in punctate cytoplasmic patterns, consistent with localization to lamellar body-like structures. LV transduced A549 ABCA3^L101P^ cells showed little overlap of ABCA3:GFP with mutant ABCA3^L101P^:mCherry, consistent with the known mistrafficking and ER retention associated with the L101P mutant. The ABCA3:mCherry phenotype in the ABCA3^L101P^ cells was mostly diffuse, however, some punctate structures were visible at higher magnification. Type 2 ABCA3 mutants E292V and E690K are expected to traffic normally to lamellar body-like organelles but have impaired phospholipid transport. As hypothesized, LV-mediated ABCA3 gene complementation of E292V and E690K cells demonstrated co-localization of GFP and mCherry (yellow, **Fig. 3A**). Of note, the diameter of the mCherry^+^/GFP^+^ vesicles increased significantly in each of these mutant lines following transduction with LV-ABCA3:GFP (**Fig. 3B**).

**Figure 3:**
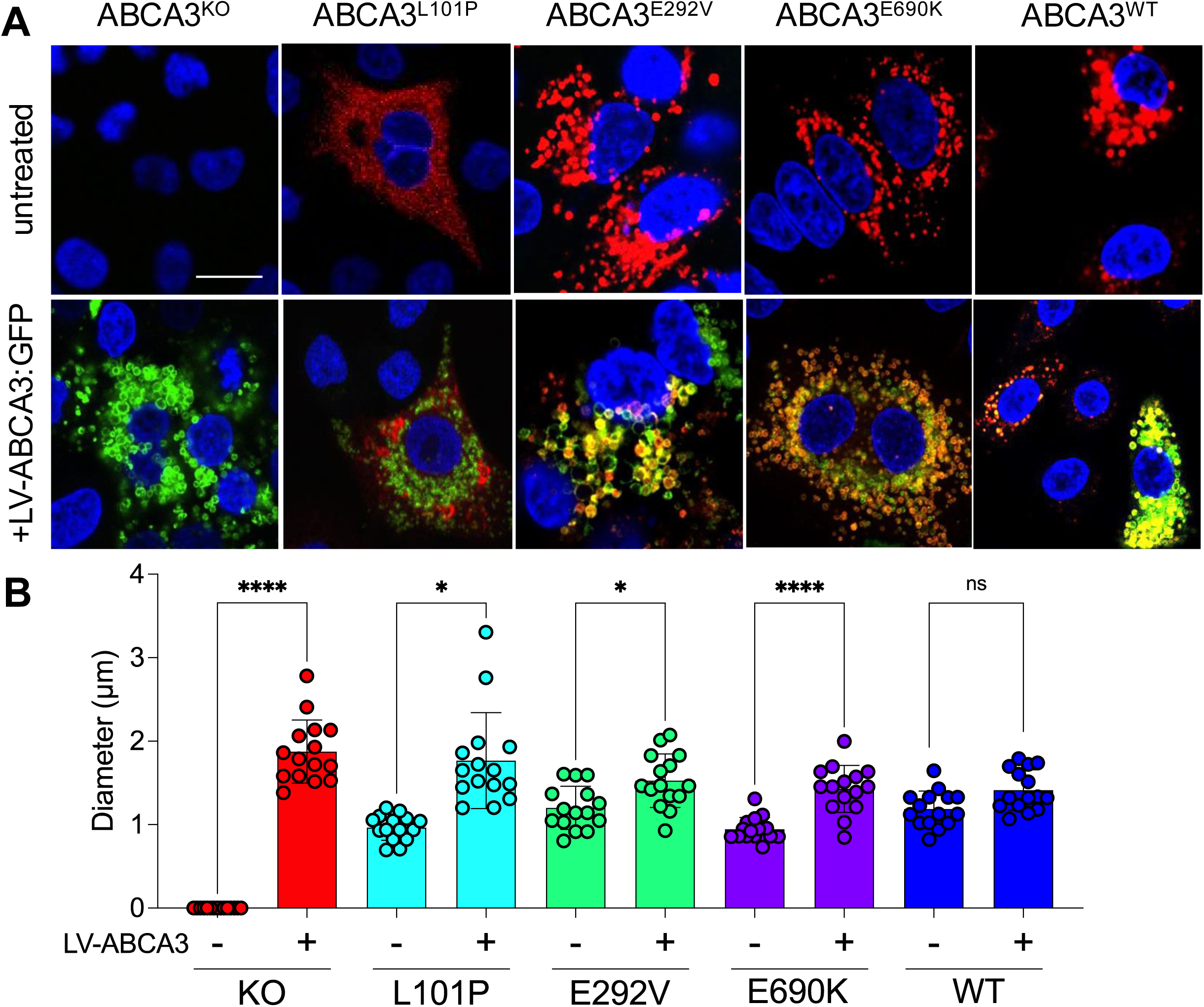
LV-ABCA3:GFP increases lamellar body-like structure size in all variant cell lines. A) Confocal microscopy images of A549 cell lines that stably express individual pathogenic *ABCA3* variants were collected. Top panels: untransduced parental lines. ABCA3^KO^ cells do not express an mCherry tag; ABCA3^L101P,^ ABCA3^E292V^, ABCA3^E690K^, ABCA3^WT^ cell lines include the mCherry tag. Bottom panels: LV-ABCA3:GFP transduced cells. GFP tag expressed from vector. Yellow indicates co-localization between mCherry and GFP tags. All fluorescence expression is from ABCA3:mCherry or ABCA3:GFP, no immunostaining was performed. B) Quantification of diameter of lamellar body-like structures was measured using ImageJ. Parental (-) lines show diameter of mCherry^+^ vesicles; transduced (+) lines show diameter of GFP^+^ vesicles. Scale bar = 15 µm. Each data point indicates a technical replicate; *p<0.05, ****p<0.0005.

Prior to cleavage, ABCA3 is translocated to the ER following translation and routed to the Golgi^7,10,26,27^. Next, ABCA3 is cleaved at the N-terminus inside LAMP3-positive vesicles^9^. We immunostained for LAMP3/CD63 to detect vesicles in the A549 cell lines that express *ABCA3* variants. In each cell line transduced with LV-ABCA3:GFP, we observed punctate GFP expression co-localizing with LAMP3/CD63 (**Fig. 4**). As expected, no GFP was observed in untransduced cells. Using Mander’s coefficient to compare co-localization, we found that co-localization patterns of type 2 mutants (E292V, E690K) were similar to wild-type ABCA3, whereas cells that express ABCA3^L101P^ demonstrated co-localization patterns more similar to ABCA3^KO^ cells (**Fig. 4B**).

**Figure 4:**
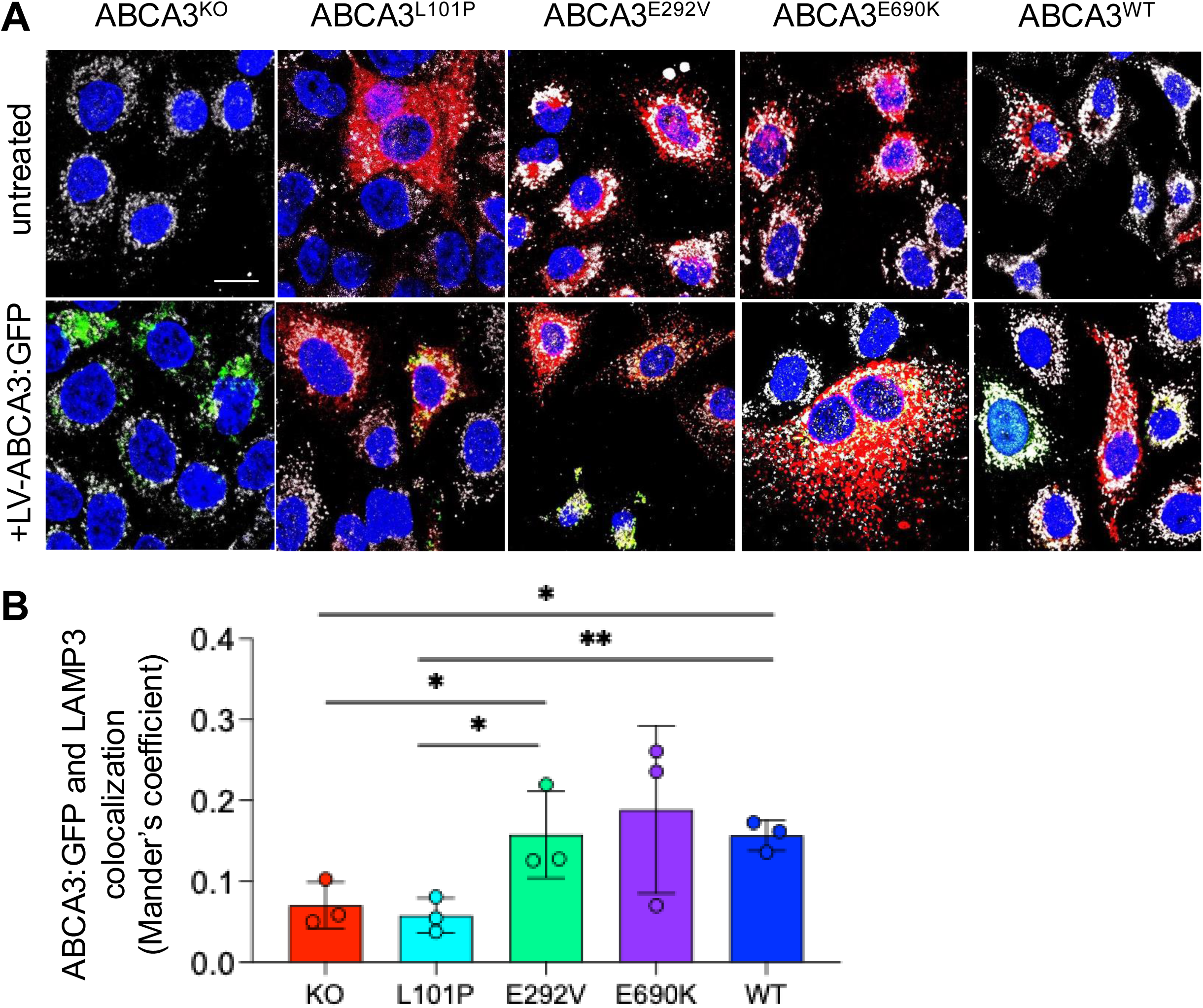
LV-ABCA3:GFP co-localizes with LAMP3. A) A549 cell lines stably expressing pathogenic *ABCA3* variants untreated or transduced with LV-ABCA3:GFP were immunostained for LAMP3 and imaged for co-localization. Blue= DAPI, green= ABCA3:GFP, red= endogenous ABCA3:mCherry, white= LAMP3. Scale bar = 15 µm. B) Mander’s coefficient showing overlap of ABCA3:GFP and LAMP3 (n=3).

In ABCA3^WT^ A549 cells, the lamellar body-like structures have wavy, non-compacted lamellae, similar to lamellar bodies in AT2 cells from healthy infants^17,19,28,29^. Using transmission electron microscopy (TEM), we examined the lamellar body-like structure morphology for each of the untreated and LV-ABCA3:GFP transduced cell lines. Lamellar body-like structure morphology for each condition was evaluated (**Supplemental Table 1**). As previously reported, we found that ABCA3^KO^ or ABCA3^L101P^ cells have electron dense and smaller lamellar body-like structures, as compared to cells that stably express the type 2 ABCA3^E292V^ variant^13,20,22^. As shown in **Fig. 5A** and **5C**, respectively, the lamellar body-like structures in untreated ABCA3^KO^ or cells that express mistrafficking (type 1) ABCA3^L101P^ were smaller and had a dense body appearance (white arrows), with some small lamellar body-like structures with at least partial lamellae in ABCA3^L101P^ (white arrowheads), while those in ABCA3^WT^ cells had many well-developed, larger normal appearing lamellar-body like structures with concentric, wavy lamellae (asterisks) (**Fig**. **5I**). The lamellar body-like structures from cells that express ABCA3^E292V^ or ABCA3^E690K^ (**Fig. 5E, 5G**) had a mixture of dense bodies (white arrowheads) and structures with wavy lamellae but smaller than seen for wild type (white arrowheads). Importantly, LV-ABCA3:GFP complementation rescued, at least partially, some lamellar body-like morphology with structures with less tightly compact lamellae observed in all LV-ABCA3 transduced conditions (**Fig. 5B, D, F, H, J**). In ABCA3^KO^ cells, all lamellar body-like structures were small and electron dense with minimal tightly compacted lamellae (**Fig. 5A**). With the complementation of LV-ABCA3:GFP, we observed some lamellar body-like structures with less tightly compact lamellae (**Fig. 5B**). For all *ABCA3* variant cell lines (ABCA3^L101P^, ABCA3^E292V^, ABCA3^690K^), LV-ABCA3:GFP transduced cells had some lamellar body-like structures with more normal appearance but also some less developed lamellar body-like structures were also present, especially for ABCA3^L101P^ (**Fig.5D**). In summary, LV-mediated complementation of ABCA3 resulted in the qualitative appearance of lamellar body-like structures with less tightly compact lamellae and even some normal appearing lamellar body-like structures in the cell lines that stably express *ABCA3* variants.

**Figure 5:**
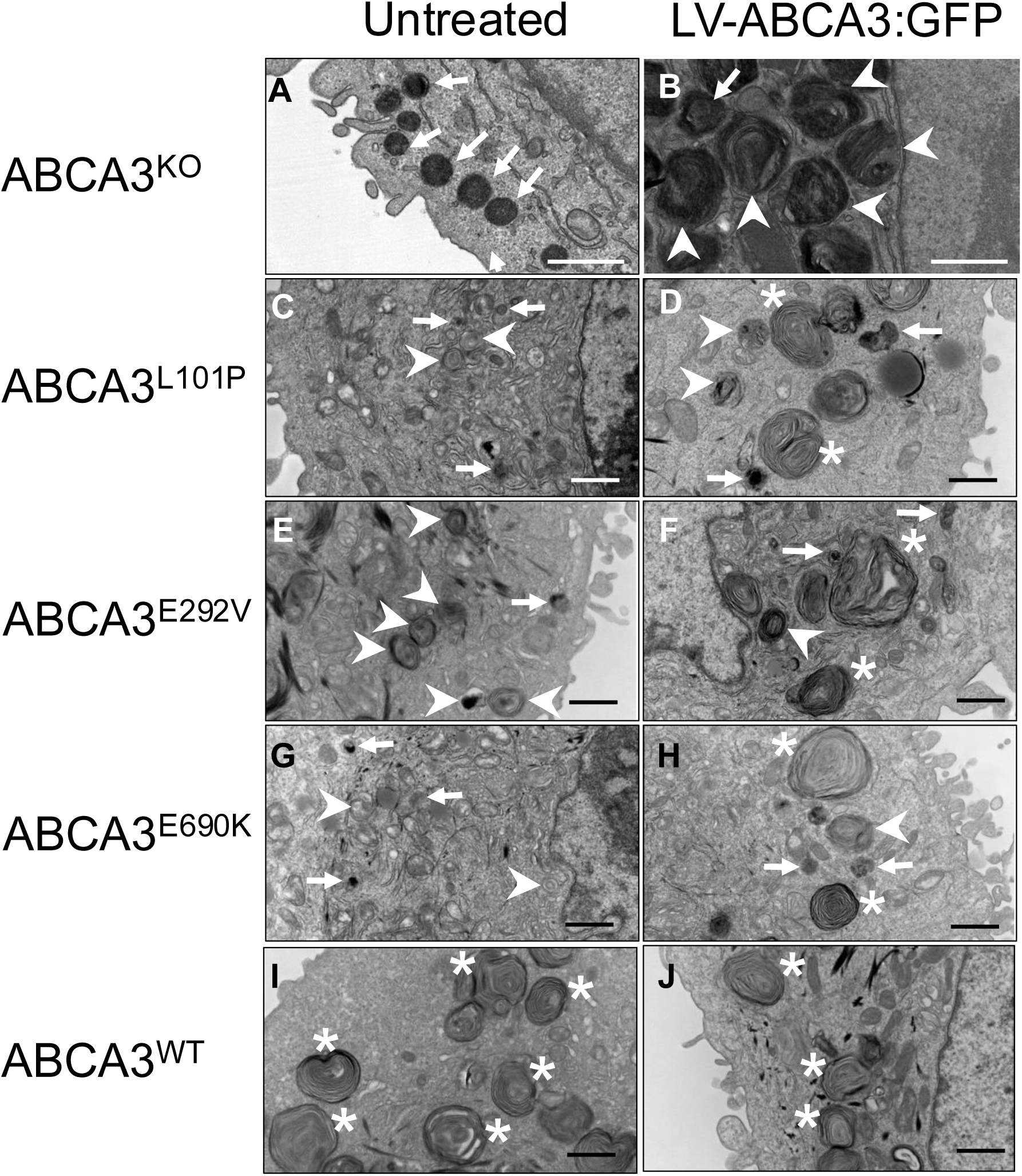
Transmission Electron Microscopy (TEM) images of A549 cell lines that stably express pathogenic ABCA3 variants. A) ABCA3^KO^, ABCA3^L101P^, ABCA3^E292V^, ABCA3^E690K^, and ABCA3^WT^ cell lines were compared to LV ABCA3:GFP transduced cells of each line. All images were taken at 20,000x. White arrows indicate representative lamellar body-like structures consistent with dense bodies, arrowheads indicate representative lamellar body-like structures with varying proportion of dense areas and wavy lamellae, and asterisks indicate representative larger well formed, normal appearing lamellar body-like structures. Of note, the TEM images from ABCA3^KO^ (untreated and LV transduced) were performed at a different facility than the TEM images from ABCA3^L101P^, ABCA3^E292V^, ABCA3^E690K^, and ABCA3^WT^ and have a two-fold final enlargement compared to other images in Figure 5. Scale bar = 1 μm.

Two distinct features of mutant ABCA3 iAT2s reported by Sun et al include a defect in cell proliferation and elevated NF-κB activation^20^. Using the A549 cells that stably express ABCA3 variants, we asked if LV-ABCA3 complementation could rescue the cell proliferation defect and decrease NF-κB activation. Cell proliferation was assayed by Edu incorporation and quantified by flow cytometry. LV complemented cells (+) exhibited increased EdU incorporation compared to untreated parental cells (-) for all *ABCA3* variants as well as WT, consistent with increased cell cycling (**Fig. 6A**). Lastly, we measured NF-κB activation using a reporter assay. Each cell line was transduced with a LV vector encoding a p50/p65 heterodimer consensus binding sequence (labeled as “NF-κB” in **Fig. 6B**), which activates luciferase in the presence of canonical NF-κB signaling. In the ABCA3^KO^ and ABCA3^L101P^ cell lines, we observed no difference in luciferase activity between +LV-ABCA3:GFP and untreated cells, but the E292V, E690K, and WT cell lines with ABCA3 complementation showed reduced luciferase activity (**Fig. 6C**). These data suggest that LV-ABCA3 rescues cell proliferation across all ABCA3 mutants studied but complementation of only type 2 mutants results in decreased NF-κB activity.

**Figure 6:**
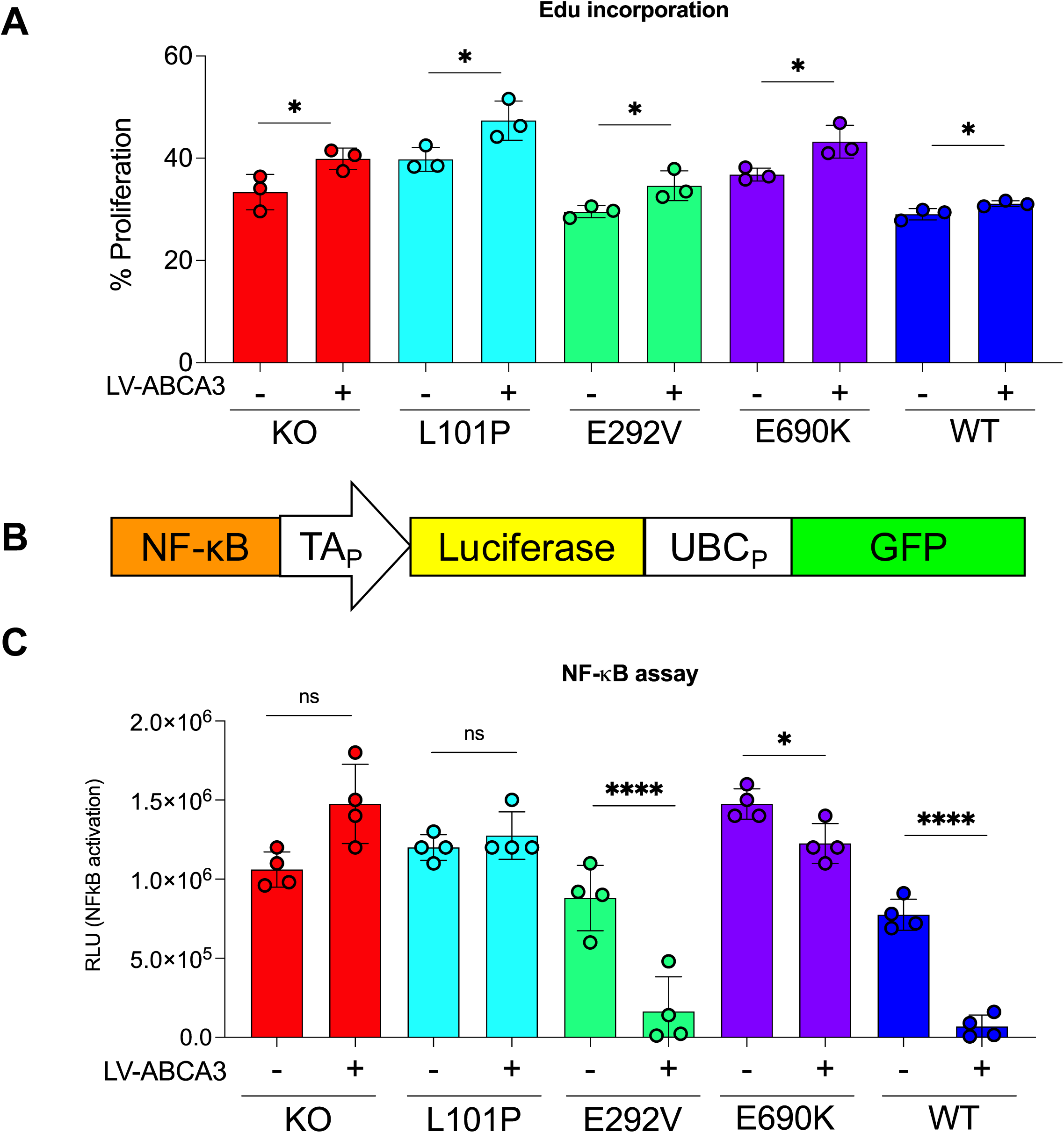
LV-ABCA3 complementation restores proliferation defect and decreases NF-κB activity. A) A549 cell lines that stably express pathogenic *ABCA3* variants and KO cells were seeded at equal densities and the next day were incubated with Click-IT Edu Pacific Blue. Cells were assayed for Pacific Blue expression by flow cytometry to measure Edu incorporation. B) Lentiviral cassette for NF-κB p50/p65 heterodimer binding site. In the presence of NF-κB, luciferase is expressed. The TAp promoter drives transgene expression when activated by NF-κB, UBCp promoter drives GFP. C) Cells were seeded and transduced with LV NF-κB-luciferase-GFP. 2 days later, cells were harvested to assess luciferase activity. N=3 or 4, *p<0.05, ****p<0.0005.

## DISCUSSION

Here we investigated the impact of LV-mediated complementation of *ABCA3* in A549 cell lines that express individual pathogenic *ABCA3* variants. We observed increases in the diameter of ABCA3+ vesicles, co-localization to LAMP3^+^ vesicles, and restoration of protein processing. ABCA3 gene complementation rescued, to some degree, the phenotypic characteristics of lamellar body-like structures in the mutant lines toward more well-developed, normal appearing lamellar body-like structures. Results from EdU incorporation suggest a proliferative advantage for LV complemented cells that express pathogenic *ABCA3* variants, consistent with previous reports^20,24^. The failure to rescue the pro-inflammatory phenotype in ABCA3^KO^ cells or ABCA3^L101P^ cells could indicate that gene addition restores phospholipid transport but does not ameliorate null or mistrafficking phenotypes. Taken together, we found that LV-ABCA3 complementation rescues several phenotypes of ABCA3 deficiency including ABCA3+ vesicle diameter, protein processing, morphology of lamellar body-like structures, cell proliferation defects, and NF-κB activity. Importantly, we found that the degree of rescue depends on the specific pathogenic *ABCA3* variant.

The ultra-rare frequency of rare and private *ABCA3* variants creates challenges for the development of variant-specific targeted treatments. A gene addition approach could be applicable to multiple pathogenic *ABCA3* variants to restore lung function or serve as a bridge to lung transplantation in the setting of progressive respiratory failure. Other therapeutic approaches being explored include small molecule studies modeled from the CFTR/ABCC7 modulator studies^30,31^. For example, in past reports, repurposed drugs resulted in increased ABCA3 protein expression and improved localization in the presence of correctors for several *ABCA3* variants in A549 cell models^17,32,33^. Importantly, the models and correction metrics described in this study could be useful for additional future small molecule screens. The development of patient-derived iAT2 cell models permits the study of additional therapeutic approaches for ABCA3 deficiency including gene editing, which will further the understanding of ABCA3 biology and expand treatment options for ABCA3 deficiency.

The integrating properties of LV vectors offer the potential for lifelong correction if progenitor cells are targeted. LV vectors have a packaging capacity of 7.5 kb, easily accommodating the *ABCA3* cDNA (∼5.2 kb). Lentiviral vectors are the leading platform for *ex vivo* gene modification of primary cells, with FDA approval of 5 CAR T-cell therapies and gene therapies for β-thalassemia (Zynteglo), cerebral adrenoleukodystrophy (Skysona), sickle cell disease (Lyfgenia), metachromatic leukodystrophy (Lenmeldy), and many additional clinical trials underway (clinicaltrials.gov). Additionally, Boehringer Ingelheim has initiated a clinical trial in the United Kingdom using a LV vector for cystic fibrosis^34,35^. These studies and numerous preclinical studies suggest that lentiviral vectors have an excellent safety profile in hundreds of patients and include some individuals with more than 10 years of follow up^36,37^. Since AT2 cells are the progenitor cells of the alveolar epithelium^38^, a lentiviral gene therapy treatment could lead to an increase in number of corrected cells over time. Rindler et al demonstrated a proliferative advantage to “corrected” *ABCA3* cells in a mouse model for ABCA3 deficiency^24^.

ABCA3 mistrafficking (type 1) and phospholipid transport mutants (type 2) are mechanistically distinct with different impacts on cell physiology, and thus may respond differently to *ABCA3* gene complementation. For example, the combined cell stress of a mistrafficking (type 1) ABCA3 mutant with reconstituted wild-type ABCA3 expression could impair fitness^39^ or reduce progenitor capacity. Here we complemented cell models representing both variant types. It is possible that regardless of the mutant type, complementing *ABCA3* would restore proliferation capacity and reduce inflammation. Alternatively, *ABCA3* gene complementation of a type 1 mutant might resolve some but not all of the disease phenotypes, while complementation of a type 2 mutant may fully restore functional readouts. The bulk mRNA sequencing data indicated an increase in cell cycling transcriptomic profiles that we confirmed by an Edu incorporation assay. These results align with previous evidence that corrected ABCA3 may confer a selective advantage for cellular division^20,24^. We found that NF-κB activity was decreased in type 2 mutants E292V and E690K complemented with LV-ABCA3, however KO and L101P cells still had elevated levels of pro-inflammatory transcripts after LV-ABCA3 transduction (**Fig 6**). This evidence suggests that *ABCA3* complementation may not rescue all ABCA3 disease manifestations equivalently.

Advantages of the A549 ABCA3 cell model include their relative ease of modification to study patient-relevant, pathogenic *ABCA3* variants and reproducibility among experiments. Although the phenotypes and transcriptomic disease signatures mirror iAT2 models, A549 cells do not express surfactant proteins B or C and therefore do not produce or release surfactant.

Therefore, testing the impact of LV-ABCA3 gene transfer on surfactant phospholipid production in iAT2 cells derived from infants and children with ABCA3 deficiency and in mouse models of ABCA3 deficiency will be important next steps. A limitation of this study is that it does not allow us to study phenotypes of type 1/type 2 compound heterozygous variants. Landing pad cells are most conducive to studying individual *ABCA3* variants rather than compound heterozygous genotypes. Developing cell models that could express compound heterozygous *ABCA3* variants will also be an important step for future personalized therapies.

In summary, LV-ABCA3 gene addition rescues several pathogenic cellular phenotypic characteristics in A549 cells that are disrupted by expression of *ABCA3* pathogenic variants. Our results suggest that while ABCA3+ vesicle diameter, co-localization, protein processing, lamellar body-like morphology, cell proliferation, and aberrant inflammatory signaling are partially rescued through LV-mediated *ABCA3* complementation, not all hallmark phenotypes are corrected similarly in the different cell lines. Additionally, investigating type 1 and type 2 ABCA3 mutants following gene complementation allows us to identify variant-specific responses to gene therapy. Understanding the efficacy of *ABCA3* functional restoration in models of alveolar type 2 cells will advance gene therapy for ABCA3 deficiency.

## MATERIALS AND METHODS

### ABCA3 variant A549 cell models

The “landing pad” cells were generated as previously published by Wambach et al^17^. Briefly, A549 cells were electroporated with CRISPR/Cas9 and guide RNA targeting exon 5 of *ABCA3* to induce a frameshift variant, producing A549 *ABCA3^KO^* cells. To generate specific ABCA3 mutant cell lines, edited cells were transduced with VSVG HIV-CMV-LoxFAS-GFP-LoxP (MOI=5). Polyclonal populations were selected through dilutional cloning and inverse PCR and the lentiviral “landing pad” integration was mapped to an intergenic region as a single insertion in chromosome 12. Next, a Cre-recombinase expression plasmid and variants of *ABCA3* cassette of LoxFAS-CMV-*ABCA3*:mCherry-LoxP were co-transfected into the landing pad cells. Cells were then FACS sorted for mCherry^+^/GFP^-^ cell populations to generate A549 cell lines that stably express individual *ABCA3* variants.

### Viral vector packaging

*ABCA3* gene augmentation studies employed a previously published lentiviral vector (pHAGE-EF1aL-WT_ABCA3:GFP; Addgene #188548^20^) encoding an *ABCA3:GFP* cDNA which expresses a single polypeptide to generate a fusion protein. This cassette is driven by a constitutively and ubiquitously active EF1αL promoter resulting in over-expression of the ABCA3:GFP fusion protein in A549 cells (termed LV-ABCA3:GFP). This self-inactivating (SIN) vector is in the pHAGE backbone and includes the Woodchuck hepatitis virus Post-transcriptional Regulatory Element (WPRE) after the coding sequence. The vector was produced at the University of Iowa Viral Vector Core (https://www.medicine.uiowa.edu/vectorcore) using a four-plasmid transfection method as previously described^40,41^. Vectors were titered using droplet digital PCR^42^ and/or by flow cytometry. Lentiviral titers ranged from 10^8^-10^9^ transducing units/ml.

### Transduction and FACS

A549 cell lines that stably express individual *ABCA3* variants were seeded in 24 well plates and transduced at MOI=20 overnight. 72 hours post-transduction, cells were lifted in TrypLE and GFP^+^ cells were sorted by fluorescence-activated cell sorting (Fusion, BD). Cell populations were gated loosely to ensure punctate GFP collection and repeated a second time to ensure a pure population of cells. GFP^+^ cells were cultured for 48 hours post-sorting and expanded to perform the experiments presented. Each figure is the result of 4 technical replicates.

### RNA and protein isolation

RNA was isolated using the Zymo RNAeasy Kit (Qiagen, Hilden, Germany). Protein was collected by seeding cells at 50,000 cells per well in a 24-well plate and harvested the following day. For each sample, four wells were harvested. Proteins were extracted using RIPA buffer (R0278, Sigma-Aldrich, Burlington, Massachusetts), supplemented with a cOmplete, Mini, EDTA-free Protease Inhibitor Cocktail (11836170001, Sigma-Aldrich, Burlington, Massachusetts) to ensure protein integrity. Protein concentration was quantified using a BCA assay prior to performing western blot analysis.

### qRT-PCR

For each sample, 0.25 µg of total RNA was used as the template and brought to a total volume of 10 µl with nuclease-free H₂O. Using a High Capacity cDNA Reverse Transcription Kit (4368814, Thermo Fischer Scientific, Waltham, Massachusetts) a 2X reverse transcription (RT) master mix was added to the RNA sample in a volume of 10 µl, resulting in a final reaction volume of 20 µl per tube. The samples were incubated in a thermocycler for cDNA synthesis following standard RT protocols. Quantitative PCR (qPCR) was performed using a 384-well plate, with a total reaction volume of 10 µl per well. The qPCR reaction master mix included the following components: *Power*SYBR Green PCR Master Mix (A25742, Thermo Fischer Scientific, Waltham, Massachusetts) at 2X, forward primer (10 µM), reverse primer (10 µM), 100 ng of cDNA, and nuclease-free water to bring the final volume to 10 µl. Primer sequences include ABCA3: Fwd: 5’GGCCATCATCATCACCTCCCACAGCA 3’, Rev: 5’AGCGCCTCCTGTTGC CCTTCACTCTG 3’, GAPDH: Fwd: 5’GTCATCCCTGAGCTGAACG 3’, Rev: 5’ CTCCTTGGA GGCCATGTG 3’. The qPCR reactions were run by the Iowa Institute of Human Genetics at the University of Iowa for amplification. Fold change (2^-²²Ct^) was calculated using GAPDH as a housekeeping gene. All reactions were performed in quadruplicate for each sample to ensure accuracy and reproducibility.

### Western blotting

Total protein quantitation was determined using a BCA Protein Assay kit (Pierce, Rockford, IL). Samples containing 50 µg total protein were prepared under reducing conditions and denatured at 37°C for 30 minutes. The proteins were separated on a 3-8% Tris-Acetate gel (Criterion XT Precast Gel) and transferred onto PVDF membranes. Membranes were blocked with 5% milk/TBST for 1 hour followed with an overnight incubation at 4°C in a rabbit anti-human ABCA3 antibody (1:100 dilution, custom antibody produced by Genemed Synthesis, Inc., San Antonio, TX). Membranes were washed and incubated for 1 hour in horseradish conjugated anti-rabbit secondary antibody (1:50,000 Goat anti-rabbit IgG HRP, Invitrogen). Protein signal was detected using an enhanced chemiluminescent substrate (SuperSignal West Femto, Pierce, Rockford, IL). Membranes were then stripped for 30 minutes at room temperature in Western Stripping Buffer (Restore Western Blot Stripping Buffer, Thermo Fisher Scientific, Waltham, MA), washed, and incubated with a rabbit anti-vinculin antibody (1:1000 dilution, Invitrogen) for 2 hours. Membranes were then washed and incubated for 1 hour in horseradish conjugated anti-rabbit secondary antibody (1:20,000 Goat anti-rabbit IgG HRP, Invitrogen, Waltham, MA). Protein signal for vinculin was detected using an enhanced chemiluminescent substrate (SuperSignal West Pico, Pierce, Rockford, IL). Protein expression was read using an Odyssey M Imaging System.

### Immunostaining

Cells were seeded onto 4 well chamber slides at a density of 40,000 cells per well. The cells were cultured in a humidified incubator at 37°C for 48 hours to allow for optimal growth and adherence. After 48 hours of incubation, the cells were fixed using 4% paraformaldehyde (PFA) in phosphate-buffered saline (PBS) for 15 minutes at room temperature. Following fixation, the cells were washed three times with PBS to remove excess PFA. The fixed cells were then maintained in PBS until further processing. To facilitate antibody access, the cells were permeabilized using a solution of 0.3% Triton X-100 in PBS for 15 minutes at room temperature. Blocking was performed to reduce non-specific binding of antibodies. The cells were incubated in Superblock for 45 minutes at room temperature. The cells were incubated with primary antibodies diluted at 1:200 and 1:500 in Superblock overnight at 4°C. The primary antibodies LAMP3 (12632-1-AP, ProteinTech, Rosemont, IL), were used at 1:200. The cells were then treated with a far-red fluorescent secondary antibody diluted at 1:600 in Superblock for 1 hour at room temperature in the dark. The slides were mounted with Vectashield containing DAPI to stain the nuclei, and fluorescent imaging was performed using a confocal or fluorescence microscope to visualize the staining results.

### Transmission electron microscopy (TEM) and lamellar body formation classification

To analyze the effects of different genetic variants on cell morphology and function, A549 cell lines that stably express *ABCA3* variants: L101P E292V, E690K, and their LV transduced versions as well as *ABCA3* wild-type (WT) and a knockout (KO) lines were utilized. Each cell line was seeded at a density of 50,000 cells per well, followed by a two-day incubation period to allow for sufficient growth and attachment. After this incubation, cells were harvested and prepared for further analysis. Fixation of the samples was performed using transmission electron microscopy (TEM) fixatives to preserve cellular structures for ultrastructural examination. Samples were subsequently sent to both Mayo Clinic Microscopy and Cell Analysis Core and The University of Iowa Central Microscopy Research Core for processing and TEM imaging. This approach allowed for high-resolution imaging and comparison of cellular features across different genetic backgrounds, enabling us to investigate how specific *ABCA3* variants influence cellular architecture and potentially contribute to disease mechanisms. At least 4 images from each condition were analyzed by a pediatric pathologist (F.V.W.) at Washington University in St. Louis. Lamellar body structures were qualitatively reviewed for ultrastructural features including dense bodies, lamellae formation, looseness/waviness of lamellae appearance, and size.

### Bulk RNA sequencing and analysis

RNA Isolation was extracted from all 10 cell lines using the Zymo RNAeasy Kit (Qiagen) following the manufacturer’s protocol. All RNA samples were subsequently submitted to the University of Iowa Institute of Human Genetics for quality assessment and processed for bulk mRNA sequencing to assess transcriptomic profiles across the different cell lines. The quality of the raw data was assessed using FastQC v.0.11.7^43^. The sequence reads were aligned to the GRCh38 reference with added GFP sequence using STAR v.2.6.0^44^. Counts per gene were summarized using the featureCounts function from the subread package v.2.0.3^43^. The edgeR package v.4.2.0^45^ was used to import, organize, filter and normalize the counts and the matrix of counts per gene per sample was then analyzed using the limma/voom normalization method^45^. Genes were filtered based on the standard edgeR filtration method using the default parameters for the “filterByExpr” function. After exploratory data analysis with Principal Component Analysis (PCA), contrasts for differential expression testing were done for each of the transduced samples vs mock (controls) for the untransduced samples. Differential expression testing was also conducted to compare the gene expression between the untransduced variant lines to investigate the differences between ABCA3 mutations. The limma package v.3.60.0^46^ with its voom method, namely, linear modelling and empirical Bayes moderation was used to test differential expression (moderate t-test). P-values were adjusted for multiple testing using Benjamini-Hochberg correction (false discovery rate-adjusted p-value; FDR). Differentially expressed genes for each comparison were visualized using Glimma v.2.14.0^47^, and FDR<0.05 was set as the threshold for determining significant differential gene expression. Functional predictions were performed using the fgsea v.1.30.0 package^48^ for gene set analysis using the Molecular Signatures Database (MSigDB) Hallmark gene sets. RNA-seq data has been deposited in the GEO Database: GEO ID: GSE302633 with the secure token: qbwlasqaxjgrnsx.

### EdU incorporation

The Click-iT™ EdU Pacific Blue™ Flow Cytometry Assay Kit (C10418, Thermo Fischer Scientific, Waltham, Massachusetts) was utilized to measure EdU incorporation. Cells were seeded at a density of 40,000 cells per well in a 24-well plate and allowed to adhere overnight. The following day, cells were incubated with Click-iT EdU (10 µM final concentration) in culture media for 2 hours. 24 hours after EdU labeling, cells were harvested, washed with BSA, and subsequently fixed and permeabilized according to the protocol provided by the Click-iT EdU Flow Cytometry Assay Kit. A Click-iT reaction cocktail was prepared and added to each sample, followed by a 30-minute incubation at room temperature, protected from light. After incubation, cells were washed to remove excess reagents. Samples were then analyzed by flow cytometry to detect EdU incorporation, indicative of DNA synthesis and cell proliferation. The assay was performed in quadruplicate, and a mock control (no EdU) was included to set appropriate gating for data analysis.

### NF-κB assay

Cells were seeded at a density of 40,000 cells per well in a 24-well plate. The following day, cells were transduced with an NF-κB gene reporter expressing luciferase following NF-κB activation and UbC driving GFP expression (Addgene #49343) at a multiplicity of infection (MOI) of 50. The transduction was performed over a 2-hour incubation period, after which complete media was added to each well to cover the transduced cells. Fresh media was applied the following day for cells to incubate for two days. The NF-κB reporter assay was performed 48 hours post-transduction to measure NF-κB activity. At this time, cells were lysed using Cell Culture Lysis 5X Reagent (E153A, Promega, Madison, Wisconsin) at 100 μl and then the Luciferase Assay Substrate (E151A, Promega Madison, Wisconsin) was added at 100 μl for detection. Luminescence was detected using the SpectraMax Luminometer. Luminescence values, indicative of NF-κB activation, were recorded and analyzed to assess gene reporter activity across the experimental conditions.

## FIGURE LEGENDS

Supplement Table 1: Variation in lamellar body-like structure morphology present in TEM images

## Acknowledgements

We thank Thomas Moninger and the Central Microscopy Research Facility for performing the electron microscopy. We thank Susan Stamnes, Jeremy Coffin, and the Viral Vector Core for their assistance in lentiviral production, titration, and quality control. We thank Jennifer Bartlett and Angela Liu for their critical review of this manuscript. This work was supported by the NIH (R03TR004814, R01HL171035, P01HL152960, P30DK54759, R01HL133089, R01HL095993, R01HL149853, P01HL170952, TL1TR001410, and N01 75N92025R00004), the Cystic Fibrosis Foundation (MCCRAY25G0, SINN25G0, STOLTZ23R0).

## Itemized Funding

National Institutes of Health grant R03TR004814 (PBM, DNK)

National Institutes of Health grant R01HL171035 (PLS, PBM)

National Institutes of Health grant R01HL149853 (JAW)

National Institutes of Health grant P01HL152960 (PBM)

The University of Iowa Precision Medicine Center for Cystic Fibrosis, National Institutes of Health grant P30DK54759

National Institutes of Health grant R01HL133089 (PLS)

Children’s Discovery Institute at St. Louis Children’s Hospital/Washington University School of Medicine (JAW)

Iowa Cystic Fibrosis Foundation Research & Development Program STOLTZ23R0

American Society of Gene and Cell Therapy Career Development Award (ALC)

Cystic Fibrosis Foundation grant SINN25G0 (PLS)

Cystic Fibrosis Foundation grant MCCRAY25G0 (PBM)

Roy J. Carver Chair in Pulmonary Research (PBM)

National Institutes of Health grants R01HL095993, P01HL170952, and N01 75N92025R00004 (DNK)

National Institutes of Health grant TL1TR001410 (EH)

## Notes

### Competing Interest Statement

The authors have declared no competing interest.

