## Supplemental Table for "Lentiviral-mediated gene complementation rescues pathogenic *ABCA3* variants"

| Description | Examples |
|---|---|
| Electron dense structure with no, minimal, or very tightly compacted lamellae (dense body) | 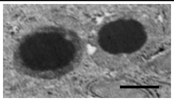 |
| Structure with varying proportions of electron dense areas and less tightly compacted lamellae, smaller than normal appearing lamellar bodies | 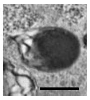 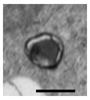 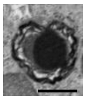 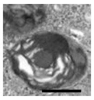 |
| Larger structure with less compact, wavy, concentric lamellae, with or without central or eccentric core matrix (similar appearance to normal lamellar body) | 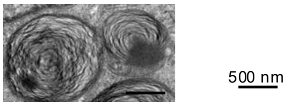  |

**Supplemental Table 1:** Variation in Lamellar Body-Like Structure Morphology Present in EM Images
