## Supplemental Figure for "Lentiviral-mediated gene complementation rescues pathogenic *ABCA3* variants"

### *Supplemental Figure 1*

**A**

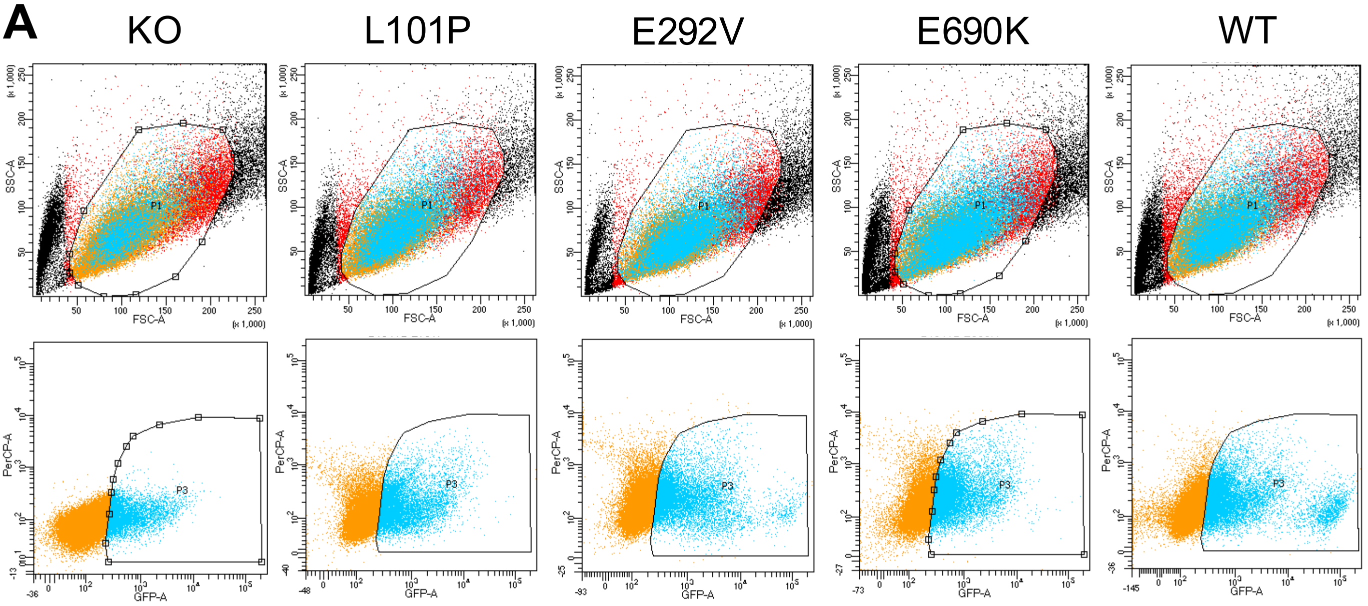


**B**

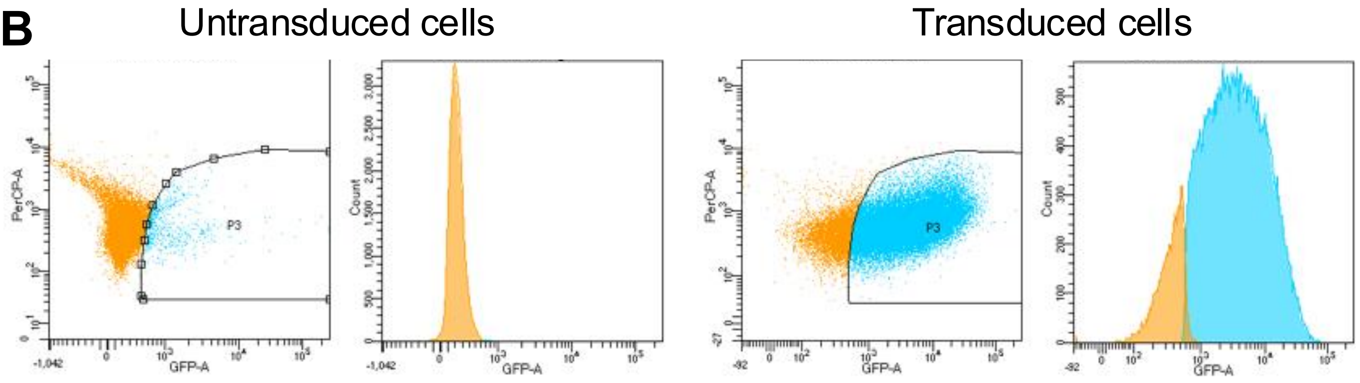


*Supplemental Figure 1: Flow cytometry gating strategy for collecting GFP positive cells.* A) Flow cytometry gating strategy for collecting GFP positive cells from ABCA3:GFP expression cassette. Top row: FSC-A vs. SSC-A gates drawn to include bulk cell population (red). Bottom row: GFP gates were drawn to collect $GFP^+$ cells transduced with LV-ABCA3:GFP (blue). Negative population (not collected) colored in orange. B) Example gating strategies for untransduced and transduced cells.
